# A neural network for online spike classification that improves decoding accuracy

**DOI:** 10.1101/722934

**Authors:** Deepa Issar, Ryan C. Williamson, Sanjeev B. Khanna, Matthew A. Smith

## Abstract

Separating neural signals from noise can improve brain-computer interface performance and stability. However, most algorithms for separating neural action potentials from noise are not suitable for use in real time and have shown mixed effects on decoding performance. With the goal of removing noise that impedes online decoding, we sought to automate the intuition of human spike-sorters to operate in real time with an easily tunable parameter governing the stringency with which spike waveforms are classified. We trained an artificial neural network with one hidden layer on neural waveforms that were hand-labeled as either spikes or noise. The network output was a likelihood metric for each waveform it classified, and we tuned the network’s stringency by varying the minimum likelihood value for a waveform to be considered a spike. Using the network’s labels to exclude noise waveforms, we decoded remembered target location during a memory-guided saccade task from electrode arrays implanted in prefrontal cortex of rhesus macaque monkeys. The network classified waveforms in real time, and its classifications were qualitatively similar to those of a human spike-sorter. Compared to decoding with threshold crossings, in most sessions we improved decoding performance by removing waveforms with low spike likelihood values. Furthermore, decoding with our network’s classifications became more beneficial as time since array implantation increased. Our classifier serves as a feasible preprocessing step, with little risk of harm, that could be applied to both offline neural data analyses and online decoding.

**New & Noteworthy:** While there are many spike-sorting methods that isolate well-defined single units, these methods typically involve human intervention and have inconsistent effects on decoding. We used human classified neural waveforms as training data to create an artificial neural network that could be tuned to separate spikes from noise that impaired decoding. We found that this network operated in real time and was suitable for both offline data processing and online decoding.

## Introduction

Brain computer interfaces (BCIs) have been used as both research tools to understand neural phenomena (Chase, Kass, and Schwartz 2012; Golub, Yu, and Chase 2015; Sadtler et al. 2014; Schafer and Moore 2011) and devices to improve patient control of prosthetics (Collinger et al. 2013; Hochberg et al. 2012; Velliste et al. 2008). BCIs interpret neural signals arising from electrodes implanted in the cortex using real time decoding algorithms; however, their performance is limited by the difficulty of isolating the activity of individual neurons from extracellular voltage signals (“spike-sorting”), the characteristics of the information present in individual neurons and multi-unit activity (their “tuning”), as well as the nuances of interactions among neurons (Averbeck, Latham, and Pouget 2006). Refining the raw data is necessary to improve BCI performance, but identifying waveforms that contain relevant information is challenging.

One common noise removal method is to use all waveforms crossing a minimum voltage threshold for decoding, where all waveforms on a particular channel are considered to be from a single neural “unit” (Christie et al. 2015; Lewicki 1998). Using threshold crossings also serves to reduce the amount of stored data. However, threshold crossings often capture noise and the summed activity of multiple nearby neurons, called multi-unit activity, on each channel (Rey, Pedreira, and Quian Quiroga 2015; Stark and Abeles 2007). Alternatively, since waveforms with the characteristic shape of an action potential are the focus of most offline neural data analyses, a different approach is to isolate well-defined spike waveforms from the threshold crossings via spike-sorting, and use only those waveforms for decoding (Lewicki 1998; Rey, Pedreira, and Quian Quiroga 2015). Unfortunately, most spike-sorting techniques are unsuitable for BCI use because they are time-intensive and require manual refinement.

Furthermore, even with the development of real time, automated spike-sorting algorithms (Chaure, Rey, and Quian Quiroga 2018; Chung et al. 2017), deciding which waveforms should be used by the decoding algorithm is challenging. There is typically no clear ground truth for what constitutes a spike waveform based solely on sparse extracellular recordings (Pedreira et al. 2012; Rey, Pedreira, and Quian Quiroga 2015; Rossant et al. 2016; Wood et al. 2004), except in the case of a few rare data sets (Anastassiou et al. 2015; Neto et al. 2016). While some studies have found spike-sorted units improved decoding performance (Kloosterman et al. 2014; Santhanam et al. 2004; Todorova et al. 2014; Ventura and Todorova 2015), others have found that using pooled spikes from threshold crossings resulted in comparable or better decoding that was more stable over time (Chestek et al. 2011; Christie et al. 2015; Dai et al. 2019; Fraser et al. 2009; Gilja et al. 2012). These conflicting results highlight the gap in our understanding of what information in neural recordings is most valuable for BCIs. Even within studies, there are inconsistent effects of spike-sorting on decoding, both between subjects and over time (Christie et al. 2015; Fraser et al. 2009). It is also difficult to compare the effects of spike-sorting on decoding performance across studies due to the inherent variability in spike-sorting techniques (Pedreira et al. 2012; Rey, Pedreira, and Quian Quiroga 2015; Rossant et al. 2016; Wood et al. 2004) and differences in brain regions, parameters being studied, and decoding algorithms that may influence which waveforms are the most decodable (Bishop et al. 2014; Chestek et al. 2011; Fraser et al. 2009; Lewicki 1998; Oby et al. 2016). Furthermore, it is unclear whether spike-sorting, where the goal is to define isolated single-units, is the best-suited approach for preprocessing data for decoding, where multi-unit activity has been shown to be sufficient in some cases (Stark and Abeles 2007).

Given spike-sorting’s history of variable effects on decoding performance, we sought to avoid explicitly sorting our data and instead developed an easily tunable spike classifier that could objectively and efficiently assign waveforms to one of two classes: spike or noise. To achieve this goal, we sought a classification approach that would (1) take advantage of the spike-sorted waveforms from previous analyses that formed a readily available, labeled training data set; (2) output a likelihood of being a spike for each waveform so we could simply adjust the minimum cutoff for binary spike classification; (3) once trained could operate in real time. We built a neural network classifier that satisfied these three criteria. To evaluate our classifier’s performance, we decoded the activity of sparse electrode arrays implanted in prefrontal cortex (PFC), and compared decoding accuracy of task condition with all threshold crossings to decoding accuracy with the waveforms selected by our classifier. We also assessed how decoding performance changed as a function of the stringency of our classifier (i.e. the minimum cutoff for spike classification) and explored how decoding accuracy with our classifier changed as a function of time since the array implantation (array implant age).

By removing waveforms that the network identified as unlikely to be spikes, we improved decoding performance for most sessions relative to decoding with all threshold crossings. Moreover, in the remaining sessions there was no substantial detriment to decoding accuracy, meaning we could apply our method with little risk of harming decoding performance. Our network classifications continued to improve decoding accuracy in sessions recorded long after array implantation, even though the overall decoding accuracy decreased and the array recordings became noisier. Thus, our real time, tunable waveform classifier demonstrates promise for long-term BCI applications and for efficient offline preprocessing.

## Materials and Methods

We trained a neural network, using a database of spike-sorted waveforms, to assign waveforms to a spike or noise class. We then tested the network’s classifications with another data set that was independent from the training set. In contrast to the offline, human supervised spike-sorting that was used to label the training data, our network did not sort waveforms into isolated single-units, but rather assigned spike or noise classifications to individual waveforms (hence its name *Not A Sorter, or NAS*). To evaluate the network’s classifications, we decoded task location using only waveforms labeled as spikes by the network and compared the accuracy to decoding accuracy using all threshold crossings. All experimental procedures were approved by the Institutional Animal Care and Use Committee of the University of Pittsburgh and were performed in accordance with the United States National Institutes of Health’s *Guidelines for the Care and Use of Laboratory Animals*.

### Neural recordings

We analyzed neural recordings from six adult male rhesus macaques (*Macaca mulatta*) which had previously been spike-sorted for ongoing experiments in the laboratory. Data from four subjects were used to train our spike-classifying neural network, and data from the remaining two subjects (Monkey Pe and Monkey Wa) were used to assess decoding accuracy using the network’s classifications. Raw recordings from both 96-electrode ‘Utah’ arrays (Blackrock Microsystems, Salt Lake City, UT) and 16-channel linear microelectrode arrays (U-Probe, Plexon, Dallas, TX) were band-pass filtered from 0.3 to 7,500 Hz, digitized at 30 kHz, and amplified by a Grapevine system (Ripple, Salt Lake City, UT). The inter-electrode distance on the ‘Utah’ arrays was 400 μm and on the linear arrays was 150 or 200 μm, beyond the range over which the two extracellular electrodes are likely to capture the waveform of a single neuron. For each recording session, a threshold (V_T_) was defined for each channel independently based on the root-mean-squared voltage of the waveforms (V_RMS_) recorded on that channel at the beginning of the session (i.e. V_T_ = V_RMS_*X, where X was a multiplier set by the experimenter). Each time the signal crossed that threshold, a 52-sample waveform segment was captured (i.e. a threshold crossing), with 15 samples prior to and 36 samples following the sample in which the threshold excursion occurred.

### Offline spike-sorting

All data in this work were spike-sorted to identify well-defined single-units for previous studies. This pre-sorted data was used to create the labeled training set for this study. Waveform segments were initially sorted into spike units and noise using a custom, offline MATLAB spike-sorting algorithm that used an automated competitive mixture decomposition method (Shoham, Fellows, and Normann 2003). These automated classifications for each recording session were subsequently refined manually by a researcher using custom MATLAB software (https://github.com/smithlabvision/spikesort). The researcher modified the classifications on each channel based on visualization of the overlaid waveform clusters, projections of the waveforms in PCA space (to assess how many clusters were present), the inter-spike interval distribution of any potential single-unit (ensuring it contained few waveforms separated by less than a sensible refractory period), and whether each potential single-unit was stationary (i.e. present for the majority of the recording session). While only a single researcher spike-sorted the waveforms from any particular session, the data in the training set (7 sessions total) collectively consisted of data spike-sorted by four different researchers. If multiple unique spike waveform shapes were present on a channel, the sorter would label those as different units.

### Training data set

We sought a diverse neural waveform training set that captured a variety of subjects, brain regions, implanted array ages (i.e. number of days since the array was implanted) and recording devices (Table 1). The four training data set subjects each had a 96-electrode ‘Utah’ array implanted in right or left hemisphere visual area V4. Two of those subjects also had 16-channel U-probe recordings from the frontal eye field (FEF), located in the anterior bank of the arcuate sulcus. Data from two sessions were used for three of the subjects: one session recorded shortly after the array implant, and one session recorded at least four months later. For the fourth subject, only a single recording session recorded shortly after the array implant was used because there were no later recordings from that array. These recordings were collected and sorted as part of previous studies which describe the experimental preparations in detail (Khanna, Snyder, and Smith 2019; Snyder et al. 2014; Snyder et al. 2015; Snyder and Smith 2015). Data in the training set were assigned a label of 0 or 1. If there were multiple single-units identified by the sorter on a particular channel, they were not distinguished in the training set and all were treated as spike waveforms (labeled as 1). We only included data from channels with very distinct spike waveforms, defined as channels with a signal-to-noise ratio (SNR) greater than 2.5 (Kelly et al. 2007). These channels still included both noise (labeled as 0) and spike waveforms (labeled as 1). The aim of using only the high SNR channels in the training set was to emphasize relatively well-isolated single-unit action potential shapes while also exposing the network to a variety of noise waveforms. Additionally, excluding low SNR channels resulted in a training set with a relatively even distribution of spike and noise waveforms (47.5% spikes, 52.5% noise).

**Table 1.**
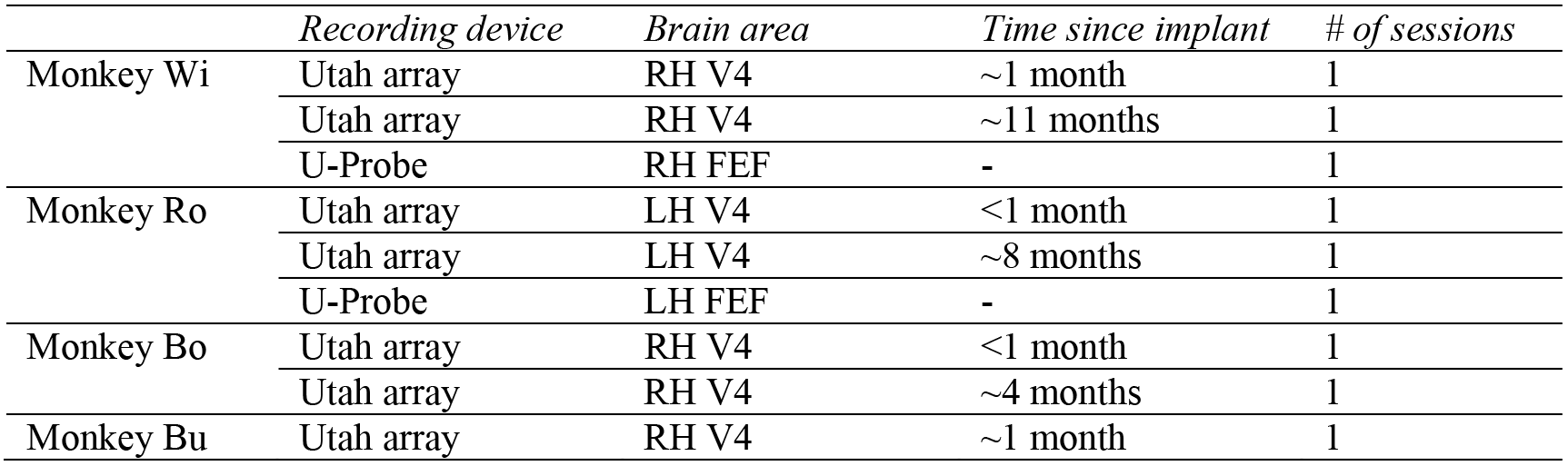
Training data set information, including time since implant (array age), brain area (RH-right hemisphere, LH-left hemisphere), and number of recording sessions. Utah arrays contained 96 channels and U-Probes had 16 channels.

|  | *Recording device* | *Brain area* | *Time since implant* | *# of sessions* |
| --- | --- | --- | --- | --- |
| Monkey Wi | Utah array | RH V4 | ~1 month | 1 |
|  | Utah array | RH V4 | ~11 months | 1 |
|  | U-Probe | RH FEF | - | 1 |
| Monkey Ro | Utah array | LH V4 | <1 month | 1 |
|  | Utah array | LH V4 | ~8 months | 1 |
|  | U-Probe | LH FEF | - | 1 |
| Monkey Bo | Utah array | RH V4 | <1 month | 1 |
|  | Utah array | RH V4 | ~4 months | 1 |
| Monkey Bu | Utah array | RH V4 | ~1 month | 1 |

Overall, the network training set consisted of 24,810,795 waveforms from four monkeys, two brain regions (V4 and FEF), two recording devices (Utah array and U-probe), and different array implant ages. These waveforms were classified as spikes or noise using the offline spike-sorting technique described above (Fig. 1a).

**Figure 1.**
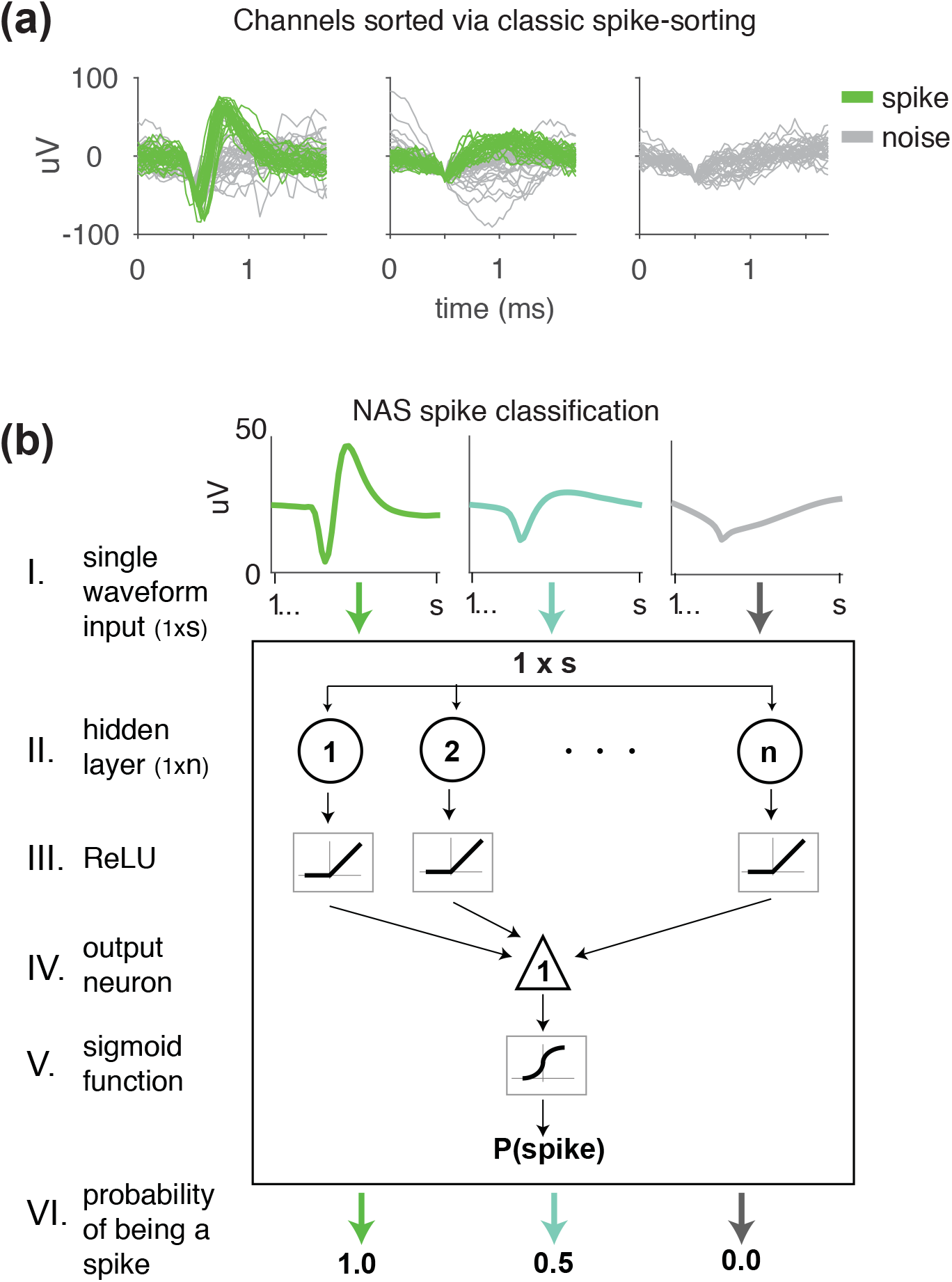
Methods of classifying waveforms: classic (manual) spike-sorting and our neural network spike classifier, Not A Sorter (NAS). (a) Spike-sorted channels with waveforms that were manually classified as spikes (green) and noise (grey). The network was trained on 24,810,795 spike-sorted neural waveforms from four monkeys. The waveforms were recorded on arrays implanted in V4 as well as U-probes placed in FEF. (b) Neural network structure and output for three sample waveforms. Each s=52 waveform input (I) was passed through a hidden layer (II) with n=50 units. The resultant linear weighting of the waveform voltages was passed through a rectified linear unit (ReLU) non-linearity (III). The output was again passed through a weighted sum (IV) followed by a sigmoid non-linearity (V). The resulting value was the network’s assessment of the likelihood that the input waveform was a spike waveform (VI). We referred to this value as P(s) or the “probability” of being a spike.

### Testing data set

We used recordings from two additional subjects, not included in the training set, for testing to ensure that the trained network was generalizable to subjects not included in its training. The two testing data set subjects (Monkey Pe and Monkey Wa) each had two 96-electrode ‘Utah’ arrays implanted, one in visual area V4 and one in dorsolateral PFC (on the prearcuate gyrus just medial to the principal sulcus, area 8Ar). Both arrays were implanted in the right hemisphere for Monkey Pe and in the left hemisphere for Monkey Wa. Only data recorded from PFC were used for the complete set of analyses (Table 2) because this provided a matched data set from two monkeys for the purpose of assessing the impact of our method on decoding and because it allowed us to test our network’s generalizability in situations where the data were recorded from a different region than the training set. However, we repeated some of the analyses in held out data from V4 and similarly found an improvement in decoding with our network (Supplementary Figure 5, https://doi.org/10.6084/m9.figshare.11808492.v1). Recordings within 50 days of the array implant were considered early sessions and recordings from over 50 days after the implant were considered late sessions (see Table 2 for session counts by subject). For Monkey Pe, the thresholds across channels were similar for all recording sessions (90% of channels had thresholds between −39 and −20 μV, median: −29 μV). For Monkey Wa, the thresholds were more permissive for the later recording sessions (> 6 months after the array implant) as the experimenters sought to extract the maximum remaining signal in arrays that were decreasing in recording quality. The median threshold of Monkey Wa’s early sessions was −32 μV (90% between −43 and −23 μV), and the median threshold of the later sessions was −15 μV (90% between −29 and −12 μV). The waveforms in the testing data set were also spike-sorted offline via the technique described above for a comparative analysis between our network classifications and offline spike-sorting.

**Table 2.** Testing data set information, including time since implant (array age), brain area (RH-right hemisphere, LH-left hemisphere), and number of recording sessions.

|  | *Recording device* | *Brain area* | *Time since implant* | *# of sessions* |
| --- | --- | --- | --- | --- |
| Monkey Pe | Utah array | RH PFC | 11-43 days | 14 |
|  |  | RH PFC | 80-147 days | 22 |
| Monkey Wa | Utah array | LH PFC | 27-40 days | 5 |
|  |  | LH PFC | 180-224 days | 11 |

### Neural Network

Using TensorFlow (Abadi et al. 2016) and Keras (Chollet 2015), we developed a neural network to classify data segments as spike or noise waveforms (Fig. 1b). The network accepts a waveform segment with ‘*s*’ samples (*s*=52), which passes through a hidden layer with ‘*n*’ units (*n*=50) that uses a rectified linear unit (ReLU) activation function. The product of the hidden layer passes through one output unit and then the network applies a sigmoid function to its output.

With the aforementioned training waveforms, the network was trained to maximize accuracy based on binary labels (0 for noise, 1 for spikes) using an Adam optimization algorithm (Kingma and Ba 2015) and a binary cross-entropy loss function in batch sizes of 100.

For a single waveform input run through the trained network, the output was a value between 0 (likely not a spike) and 1 (likely a spike), which represented the network’s assessment of the likelihood that the input waveform was a spike. While this output value was not a conventional probability, we refer to it as the probability of being a spike or P(spike) because it was a likelihood metric scaled between 0 and 1. Supplementary Figure 1 (https://doi.org/10.6084/m9.figshare.11808492.v1) provides more intuition regarding the hidden layer units and how the network assessed waveforms. In order to classify the waveforms using the network’s output probability, we set a minimum P(spike) for a waveform to be classified as a spike. We referred to this minimum probability as the *γ* threshold. A *γ* threshold of 0 classified all of the waveforms that were captured for the session as spikes; this is often referred to as ‘threshold crossings’ in the literature. Increasing *γ* resulted in fewer waveforms classified as spikes because the P(spike) cutoff was higher.

We refer to our neural network as Not A Sorter (NAS), because it classifies waveforms as spikes or noise but does not attempt to sort the spikes into separable single-units. Custom Python and MATLAB scripts used to train the neural network and classify waveforms, as well as sample training and testing data, are available at https://doi.org/10.6084/m9.figshare.11808492.v1. The NAS software is also integrated into our custom MATLAB “Spikesort” package (https://github.com/smithlabvision/spikesort).

### Memory-guided saccade task

The two testing data set subjects (Monkey Pe and Monkey Wa) performed a memory guided saccade task (Fig. 2a). Each subject fixated on a central point for 200 ms. Then, a target stimulus flashed at a set amplitude and direction for 50 ms, followed by a 500 ms delay. There were a total of 40 possible conditions: the radius could be one of five eccentricities and the target direction could be one of eight (spaced in 45° steps). After the delay, the fixation point disappeared instructing the subject to make a saccade to the location where the target flashed. For some sessions, during the saccade the target reappeared to help the subject; however, for all analyses we only used data from the beginning of the delay period (i.e. prior to the saccade). Subjects were rewarded with water or juice for making a saccade to the correctly remembered target location. For all decoding analyses, the decoded condition was the target location. Trials at a single eccentricity (Monkey Pe: 9.9°, Monkey Wa: ~7.6°) were used to maximize the number of recording sessions that could be compared. Thus, there were eight unique target locations to decode.

**Figure 2.**
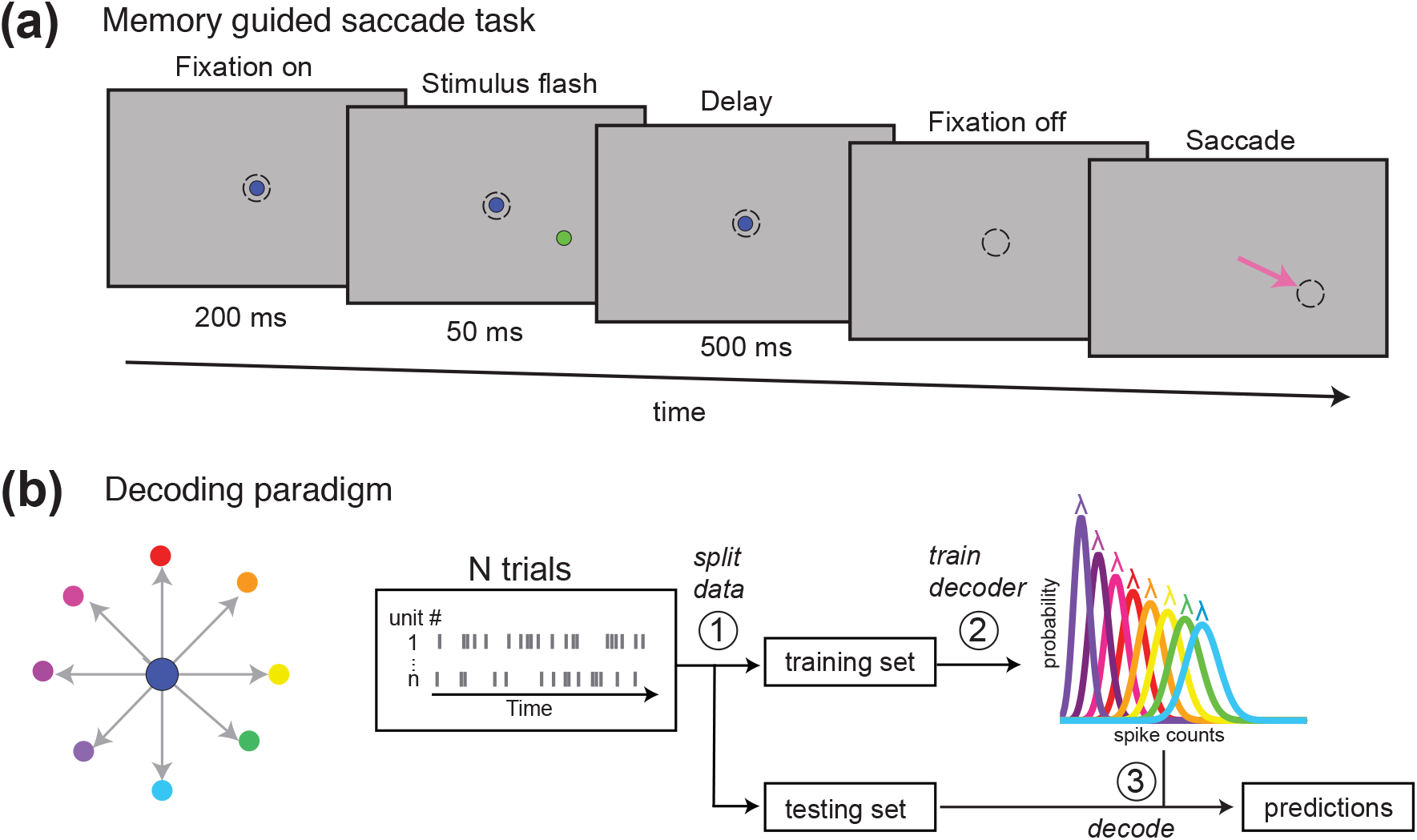
Offline decoding of planned direction from a memory guided saccade task. (a) Memory guided saccade task. The monkey fixated on a central point. A brief stimulus flash occurred followed by a delay period. When the fixation point turned off, the subject was required to make a saccade to the location of the previously flashed stimulus. (b) Decoding paradigm. The stimulus flash could occur at 8 different angles around the fixation point. A Poisson Naïve Bayes classifier was used to decode stimulus direction offline using spikes recorded from a 96-electrode Utah array in PFC 50 ms after target offset. For each recording session, we used 5-fold cross validation, where for each fold the data were split (step 1) into a training set to create model distributions for each direction (step 2) and an independent testing set to test the accuracy of the model’s predictions (step 3). Note that the curves in step (2) do not depict actual distributions and simply represent how a Poisson decoder could use spike counts from trials in the training set to distinguish between different target conditions.

Only rewarded trials were analyzed. The number of trial repeats per condition varied over sessions. Monkey Pe had an average of 43 repeats per session (range: 25-75, sd: 17) and Monkey Wa had an average of 62 repeats per session (range: 51-83, sd: 11) for each target condition.

### Decoding

To classify target direction prior to movement from spikes in PFC, we used a Poisson Naïve Bayesian decoder (Fig. 2b). All decoding analyses were performed offline. Waveforms classified as spikes from 300 ms to 500 ms after fixation (i.e. 50 ms after stimulus offset) were used for decoding, while waveforms classified as noise were discarded. Each channel was considered a single decodable unit with one exception: to calculate decoding accuracy using manual spike-sorting, if there were multiple single-units identified by the manual spike-sorting method on a channel, each single-unit on that channel remained separate for the purposes of decoding. Units with an average firing rate across trials below 0.25 spikes per second were discarded. We allocated training and test trials using two different methods. For one analysis, we assigned 80% of the trials from each session to the training set and the remaining 20% to the test set (Fig. 4a,b). For the remaining analyses, we wanted two test sets, so we used 60% of the trials for training and 20% for each of the two test sets. The two test sets allowed us to search for an optimal *γ* threshold using the first test set and cross validate this value with the second test set. Training and test data were rotated such that each trial was used for a test set only once.

Our decoding algorithm created a Poisson distribution model for each target location (*θ*) using the average spike count for each unit (*n*_*spike*_) in the training set. For each test trial, the target location with the maximum prediction probability, *P*(*θ|n*_*spike*_), was the predicted label. In equation (1), *P*(*n*_*spike*_|*θ*) was calculated using the Poisson model developed with the training trials:

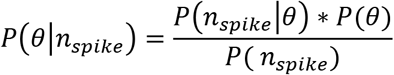

Decoding accuracy was calculated as the ratio of the number of correct prediction labels to the total number of predictions. We used 5-fold cross validation and computed the average decoding accuracy across folds.

## Results

We trained a one-layer neural network called Not a Sorter (NAS) to evaluate the likelihood that a neural waveform was a spike. We used a diverse set of waveforms in the training set from different brain regions, subjects, and array implant ages (time since array implant) in order to expose the network to a variety of waveform types. Each waveform in the training set was assigned a binary label of noise (0) or spike (1) via offline spike-sorting with manual refinement by researchers. The network was also exposed to variability in spike classification due to manual sorting because different researchers sorted different sets of waveforms in the training data. The network learned to assess how spike-like a waveform was based on binary labels, but itself output a continuous value between 0 and 1 for each waveform allowing for tunable classification. We referred to this output as the probability of being a spike or P(spike). The network classified 1000 waveforms in less than one millisecond on average (computed on a 2011 iMac with a 2.8 GHz Intel Core i7 processor). In a 10 ms bin, 1000 waveforms would be the expected output of 1000 channels each recording from a single-unit with a firing rate of 100 spikes/s. Thus, our network could be easily integrated to operate within real time computing constraints. In a realistic simulation of a real time application of our network, waveforms captured from 192 channels in a 20 ms time step could be classified in less than 0.1 ms on average, easily sufficient for the updating of a BCI cursor or other feedback.

### Qualitative assessment of NAS classifications

To assign each NAS-classified waveform a binary spike or noise label, we set a parameter called the γ threshold, which was the minimum P(spike) a waveform needed to be considered a spike waveform. We found that even a low γ threshold, such as γ=0.20 (Fig 3a), assigned most waveforms to classes that a human spike-sorter would deem appropriate. Most spike-sorters and spike-sorting algorithms search for waveforms with a canonical action potential shape – an initial voltage decrease followed by a sharp increase, a narrow peak, and a return to baseline. Increasing the γ threshold to 0.70 (i.e. only waveforms assigned a P(spike) > 0.70 were considered spikes) mimicked the effect of more selective spike-classification (Fig 3b), where the percentage of spike waveforms on the channel decreased since more waveforms were placed in the noise class and the remaining spike waveforms had a clearer single-unit shape. Considering all of the waveforms from this same channel, the range of P(spike) values coincided well with our subjective impression of the match of individual waves with a canonical action potential shape (Fig 3c), and this was also true across all channels from this array (Fig 3d). Thus, tuning the γ threshold from low values (near 0) to high values (near 1) resulted in a shift from a more permissive to a more restrictive regime.

**Figure 3.**
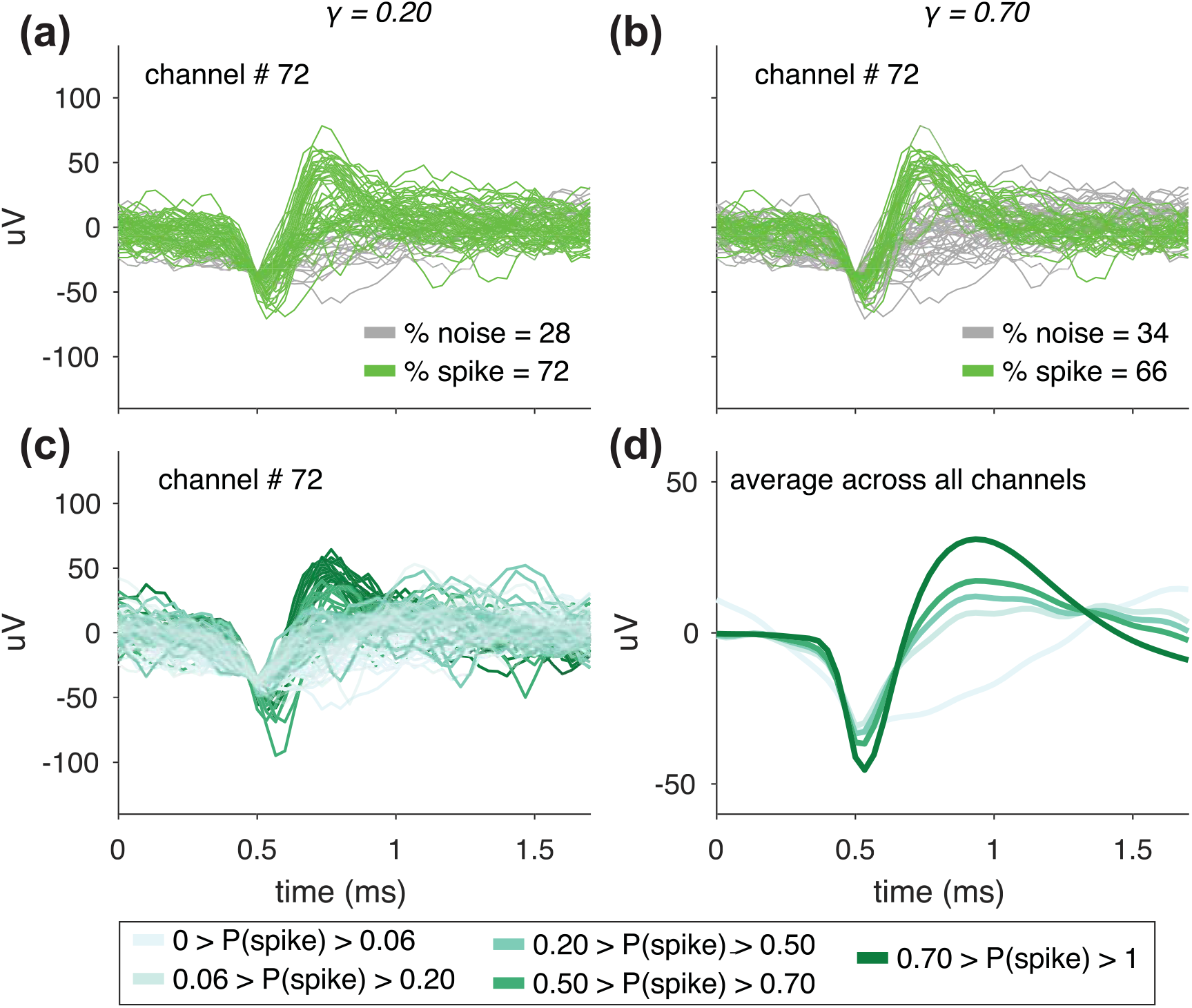
Classifying waveforms based on their probability of being a spike, P(spike), by setting γ, a tunable parameter. (a-b) Waveforms classified as spikes (green) or noise (grey) for a sample channel in Monkey Pe. The network outputs a value between 0 and 1, referred to as P(spike), for each waveform, where a value close to 1 means the network identifies that waveform as very likely to be a spike waveform. After running waveforms through the network, we set the minimum P(spike) to classify waveforms as spikes and referred to this value as the γ threshold. In (a) only waveforms assigned a spike probability greater than 0.20 (i.e. γ threshold = 0.20) and in (b) greater than 0.70 (i.e. γ threshold = 0.70) were classified as spikes. Increasing the γ threshold by definition results in a smaller percentage of waveforms captured in the spike class. (c) Waveforms from the same channel in (a) and (b) colored based on their network assigned P(spike) value. For a waveform where γ_1_ < P(spike) < γ_2_, the waveform would be classified as a spike when the threshold is γ_1_, but would be labeled as noise for γ_2_. (d) Similar to (c) except the average of waveforms across all channels within the indicated P(spike) ranges. Modifying the γ threshold tuned the stringency of the spike classifications. For the channel depicted in (a)-(c) the standard deviation of the waveform noise, computed using the method described in Kelly *et al.* (2007), was 17.1 μV.

### Quantitative assessment of NAS classifications with decoding

Although by eye the network appeared to classify waveforms reasonably well, we sought to objectively assess its performance by measuring its effect on offline decoding accuracy. Specifically, we hypothesized that decoding using spikes classified by our network would be better than using threshold crossings. We applied our network to a set of data recorded from PFC on which the network had not been trained. First, we set the γ threshold and discarded any waveforms with a P(spike) below the threshold. Next, we split the data into training and testing sets. We trained the decoder using the training set to decode the task condition (the remembered location, out of 8 possibilities) during the delay period of a memory-guided saccade task on each trial (see Methods). Then, we used the trained decoder to assess decoding accuracy in the test set(s). We repeated this process for the same data set using multiple γ thresholds between 0 (i.e., all threshold crossings were considered spikes) and 0.95 (i.e., only waveforms assigned a 0.95 or greater probability by the network were considered spikes). Since only waveforms classified as spikes were used for decoding, as the γ threshold increased, fewer waveforms remained for decoding.

Initially, we used 80% of trials to train the decoder and the remaining to test it. We analyzed cross-validated decoding accuracy as a function of γ (sample recording sessions in Fig. 4a,b). Chance decoding was 12.5% (1 out of 8) and was verified by computing decoding accuracy for shuffled test trials. The decoding accuracy using threshold crossings (γ = 0) for the sample recording sessions in Fig. 4a and 4b was close to chance levels for Monkey Pe (12.8%) and was also low for Monkey Wa (20.1%). Using the network’s classifications improved decoding accuracy relative to using threshold crossings. The peak decoding benefit of using the network’s spike classifications occurred at a low γ threshold (Monkey Pe: γ = 0.16, Monkey Wa: γ = 0.16). Being very selective about the spikes used for decoding (by setting a high γ threshold) did not have a large impact on decoding accuracy, which appeared to plateau after the peak. We found that increments of the γ threshold did not remove the same number of waveforms. A substantial proportion of waveforms were removed at the lowest γ threshold tested (especially in Monkey Pe), and small increments of the γ threshold as it approached a value of 1 could also result in large increases in the proportion of removed waveforms (especially notable in Figure 4b). For reference, when we manually spike-sorted the same data offline (see Methods), we discarded 89.2% of the waveforms in Figure 4a and 71.5% of the waveforms in Figure 4b. These values were similar to the percentage of waveforms the network removed at higher γ thresholds.

**Figure 4.**
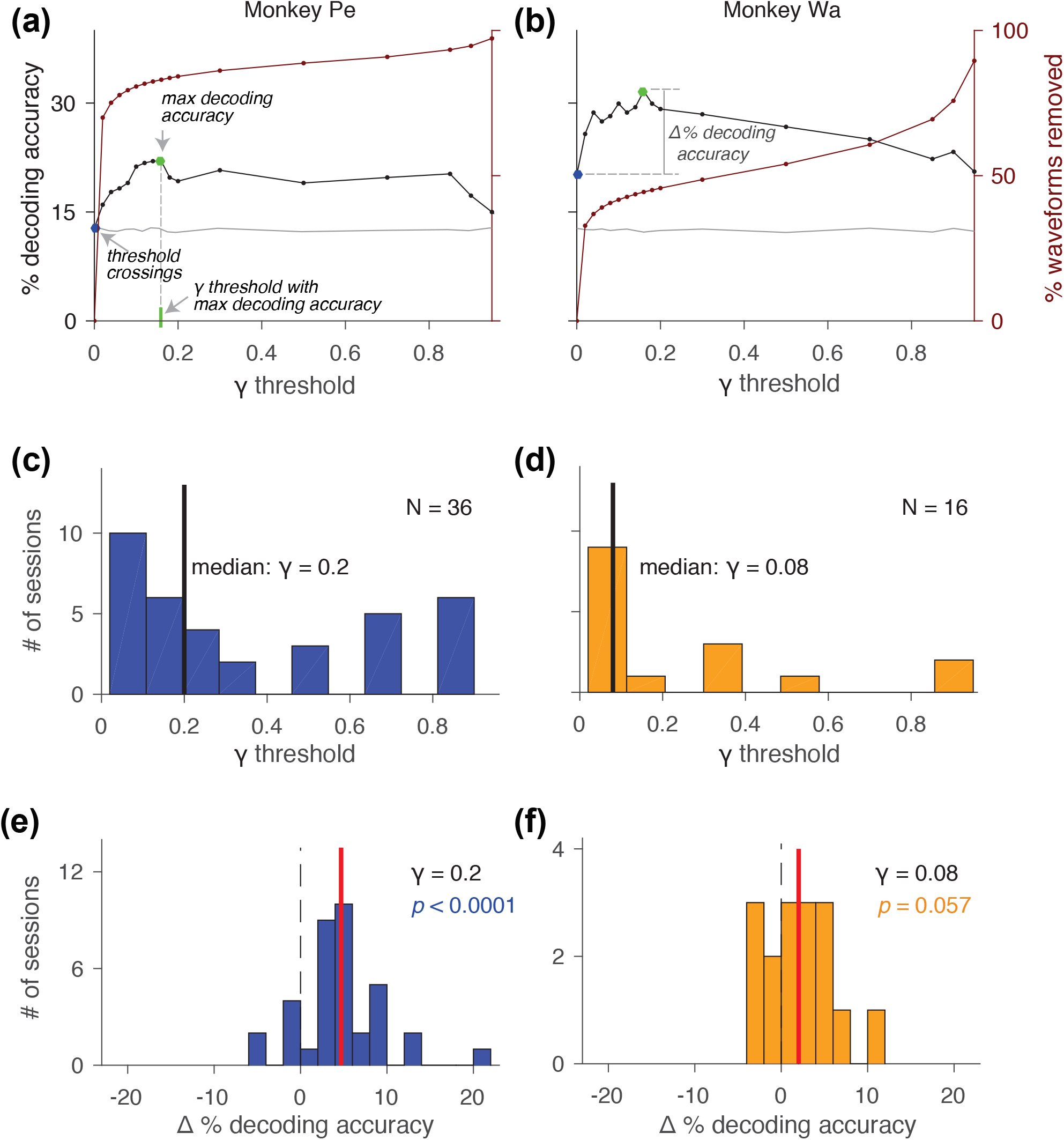
Using NAS spike classifications improved decoding accuracy in many sessions. (a-b) Percent decoding accuracy (black line) and percent of waveforms removed (maroon line) at different γ thresholds for an example recording session in Monkey Pe and Wa. Chance decoding accuracy was 12.5% (verified by computing a decoding control with shuffled test trials, grey line). The decoding accuracy increased for low γ thresholds and then reached an asymptote as γ increased. At the highest γ thresholds, decoding accuracy fell to chance (grey line). Other labeled relevant metrics include: decoding accuracy with threshold crossings (blue dot) maximum decoding accuracy (green dot), γ threshold with the maximum decoding accuracy (grey dotted line), and Δ % decoding accuracy (the difference for a given session between the decoding accuracy using the network classifications at a particular γ threshold and the decoding accuracy with threshold crossings). (c-d) Distribution of γ thresholds across sessions (Monkey Pe: N =36, Monkey Wa: N = 16) that resulted in the maximum decoding accuracy for each session. (e-f) Distribution of Δ % decoding accuracy across all sessions where the γ threshold was the median from the distributions in (c-d). Mean Δ % decoding accuracy shown in red. For both monkeys, using spikes classified by the network improved decoding accuracy on average across sessions; however, this improvement was only statistically significant for Monkey Pe (2-tailed, Wilcoxon signed rank test, Monkey Pe: p < 0.0001, Monkey Wa: p = 0.057).

Ideally, for the sake of simplicity during online BCI experiments, we could select a single γ threshold and use it for all recording sessions. We wanted to find the lowest γ threshold that improved decoding accuracy for the majority of recording sessions. We focused on lower γ thresholds because it was a conservative approach, and because our intuition from single channels and results in example sessions (Figure 4) showed it could have the greatest benefit for decoding. In order to cross-validate both our decoder and our γ threshold selection for the remaining analyses we created two test sets for decoding (training set: 60% of trials, test set 1: 20% of trials, test set 2: 20% of trials). Our general strategy was to use the first test set to identify a γ threshold that optimized decoding accuracy. We then cross-validated this selection by applying the chosen γ threshold to the data in the second test set and computing decoding accuracy for those unseen trials.

As described above, after training the decoder for each session using the training trials, we calculated decoding accuracy with the first set of test trials at each γ threshold. We found the maximum decoding accuracy at any γ threshold greater than 0, and then searched for the lowest non-zero γ threshold that resulted in a decoding accuracy within 99% of that maximum (Fig. 4c, d). Since the optimal γ threshold varied between sessions, we computed the median across sessions of these γ values and rounded it to the nearest tested value (Monkey Pe: γ = 0.2, Monkey Wa: γ = 0.08). We then used that median value as the γ threshold for all sessions in the second test set and analyzed the change in decoding accuracy from decoding using threshold crossings in that same test set, which we termed Δ % decoding accuracy (Fig. 4e, f). Combining sessions from both animals, the average improvement in decoding accuracy was 3.9% (2-tailed, Wilcoxon signed rank test; across subjects: *p* < 0.0001, Monkey Pe: *p* <0.0001, Monkey Wa: *p* = 0.057). Thus, our network’s classifications tuned to the previously described γ thresholds resulted in a net benefit for decoding performance across sessions compared to using threshold crossings. Although we could tune the γ threshold for each session to maximize the decoding, our choice of a fixed γ threshold was more consistent with the use of our network in an online decoding context, where it would be desirable to set a constant γ threshold at the start of each session rather than tuning it as a free parameter. However, an alternative strategy would be to collect a small data set at the start of each day to find the optimal γ value, and then continue experiments for the remainder of that day using the chosen value.

Given the ability of our network to improve decoding accuracy beyond that observed with threshold crossings, we took advantage of our longitudinal recordings in a fixed paradigm to understand how our network’s performance varied as a function of time. Since there is mixed evidence in the literature regarding decoding stability with long-term array implants, we were specifically interested in how the passage of time since array implant (array implant age), and concomitant degradation of recording quality, could influence our network’s performance.

### Impact of array age

First, we assessed how the overall quality of our neural data changed as a function of time. As the time since the array implant increased, the percentage of waveforms that the network assigned a very low probability of being a spike (P(spike) < 0.02) also increased (Fig 5a; Spearman’s correlation, Monkey Pe: ρ = 0.87, *p*<0.0001; Monkey Wa: ρ = 0.90, *p*<0.0001). Conversely, the percentage of waveforms assigned higher probabilities of being a spike (P(spike) > 0.70) decreased over time (Fig 5b; Monkey Pe: ρ = −0.87, *p*<0.0001; Monkey Wa: ρ = −0.94, *p*<0.0001). These classification changes were consistent with our qualitative observations that the arrays showed an increasing proportion of multi-unit activity (relative to single-unit activity) and apparent noise over time. These changes were also reflected in our manual spike-sorting (performed offline on the same data for previous studies, see Methods) where we discarded a smaller percentage of waveforms from earlier sessions that were 0-50 days post-implant (mean % waveforms removed +/− 1 sd, Monkey Pe: 36.1 +/− 8.2%; Monkey Wa: 46.1 +/− 12.2%) and a larger percentage in later sessions that were greater than 50 days post-implant (mean % waveforms removed +/− 1 sd, Monkey Pe: 87.0 +/− 7.7%; Monkey Wa: 77.8 +/− 10.3%). We further found that the ratio of the percentage of waveforms assigned a high P(spike) to the percentage of waveforms assigned a low P(spike) by the network served as a proxy for a signal to noise ratio metric, and it was highly correlated with the median signal to noise ratio (across channels) of the array in each session (Supplementary Figure 2, https://doi.org/10.6084/m9.figshare.11808492.v1). Both signal to noise metrics decreased over time, and the decoding accuracy with threshold crossings also decreased the longer the array had been implanted (Fig 5c; Monkey Pe: ρ = −0.86, *p*<0.0001; Monkey Wa: ρ = −0.69, *p*=0.004).

**Figure 5.**
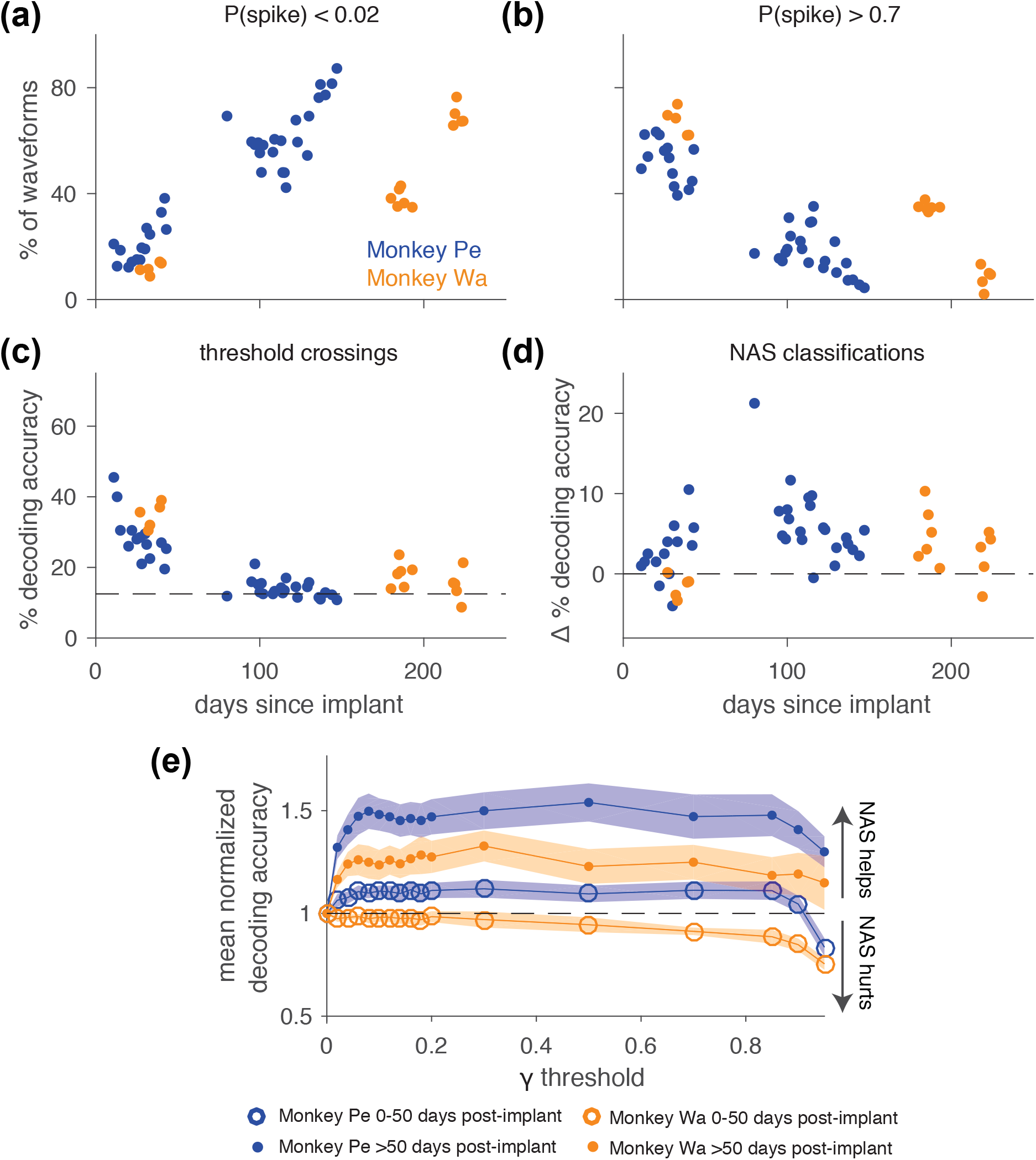
Noise on the array increased with the age of the array while decoding accuracy decreased in Monkey Pe (blue) and Monkey Wa (orange). (a) Percentage of waveforms with a P(spike) < 0.02 over time. A waveform with a P(spike) less than 0.02 was one that the network found very unlikely to be a spike. The percentage of these unlikely spike waveforms increased as the array became older (Spearman’s correlation, Monkey Pe: ϱ = 0.87, p < 0.0001; Monkey Wa: ϱ = 0.90, p < 0.0001). (b) Percentage of waveforms with a P(spike) > 0.70 over time. The percentage of waveforms that the network found to be strongly spike-like decreased as the array became older (Spearman’s correlation, Monkey Pe: ϱ = −0.87, p < 0.0001; Monkey Wa: ϱ = −0.94, p < 0.0001). (c) Decoding accuracy with threshold crossings (i.e. γ = 0) decreased as the array aged (Spearman’s correlation, Monkey Pe: ϱ = −0.86, p < 0.0001; Monkey Wa: ϱ = −0.69, p = 0.004). (d) Change in percent decoding accuracy using network-classified spikes relative to decoding accuracy with threshold crossings (Δ % decoding accuracy). We computed a distribution of the maximum γ threshold (similar to Fig. 4c-d) for sessions that were 0-50 days post array implant and used the median to set the γ threshold before computing decoding accuracy for those sessions. We repeated this for sessions more than 50 days post array implant. In Monkey Pe, using NAS classifications improved decoding accuracy (2-tailed, Wilcoxon signed rank test, p < 0.0001) and in Monkey Wa, using the classifications neither hurt nor helped decoding significantly (p = 0.09). In both subjects, the network helped decoding more in the late array sessions (>50 days post implant) compared to the early sessions (2-tailed, Wilcoxon rank sum, Monkey Pe: p = 0.01; Monkey Wa: p = 0.01). (e) Mean normalized decoding accuracy across early sessions (open circles) and late sessions (filled circles) as a function of γ threshold. Shading represents +/− 1 SEM. Using the network classifications at any γ threshold greater than zero was more helpful for late sessions than it was for early sessions.

Given the change in distribution of the types of waveforms present during the session over time, we tried to optimize the γ threshold based on time since array implant. We used the same method as in Fig. 4c-d to compute the γ thresholds with approximately maximum decoding (within 1%), except we calculated the median of the early sessions (recorded 0-50 days post implant) and the median of the late sessions (recorded >50 days post implant) separately. This did not prove to be a useful optimization as the median values were similar between early and late sessions in both Monkey Pe (γ_early_ = 0.25, γ_late_ = 0.18) and Monkey Wa (γ_early_ = 0.06, γ_late_ = 0.08). Decoding accuracy compared to threshold crossings (Δ % decoding accuracy) was significantly increased in Monkey Pe (Figure 5d, 2-tailed, Wilcoxon signed rank test, *p* < 0.0001) and was not significantly helped or hurt in Monkey Wa (p = 0.09). In line with the trends in signal and noise over time, the decoding accuracies from later sessions (> 50 days post implant) were helped more by the network classifications in both subjects than those from earlier sessions (Figure 5d, Wilcoxon rank sum, Monkey Pe: *p* = 0.01; Monkey Wa: *p* = 0.01). When we normalized the decoding accuracy at each γ threshold by the decoding accuracy using threshold crossings for each session and separately averaged across early and late sessions, the increased benefit of using the network in the later sessions compared to the earlier sessions was clear (Figure 5e).

Altogether, these results provide additional evidence that factors such as time since the array was implanted and signal to noise ratio influence decoding accuracy and affect how noise removal impacts decoding performance. Despite decreasing signal quality and increasing noise waveforms, the effect of using our network classifications for noise removal prior to decoding was consistent in that it was most often beneficial for decoding and at worst minimally detrimental. We confirmed that these results were not affected by poor γ threshold selection by using the maximum decoding accuracy regardless of γ threshold to calculate Δ % decoding accuracy (Supplementary Figure 3a, https://doi.org/10.6084/m9.figshare.11808492.v1). Our results were also not substantially influenced by variability in the number of trials across sessions (Supplementary Figure 3b, https://doi.org/10.6084/m9.figshare.11808492.v1). An even simpler alternative to our network might be to adjust the voltage threshold for capturing waveforms, to make it more or less permissive. We found that increasing this threshold (making it more negative and therefore excluding the waveforms that did not exceed it) did not improve decoding performance (Supplementary Figure 4, https://doi.org/10.6084/m9.figshare.11808492.v1), indicating that our network was not merely acting to exclude small amplitude waveforms. Lastly, we confirmed that our network was beneficial for decoding in data recorded from another brain region (V4) with different Utah arrays in the same sessions from the same animals in the test data set (Supplementary Figure 5, https://doi.org/10.6084/m9.figshare.11808492.v1).

### Comparison to spike-sorting

In light of the inconsistent effect of spike-sorting on decoding performance in the literature (Christie et al. 2015; Dai et al. 2019; Fraser et al. 2009; Todorova et al. 2014), we sought to evaluate the relative merits of manual spike-sorting and our network classifier on decoding accuracy in our data. Our goal was to place our new method (using our network to remove noise but not sort the data) in context of previous work that evaluated the impact of human supervised spike-sorting on decoding. We took advantage of the offline spike-sorting that had already been applied to these data and calculated the decoding accuracy of spike-sorted data (Fig. 6, data aggregated across subjects in the marginal histograms). Decoding using spike-sorted waveforms was better than using threshold crossings (Fig. 6 right marginal; 2-tailed, Wilcoxon signed rank test, *p* < 0.0001) similar to decoding with our network classifications (Fig. 6 top marginal, data from Fig. 5d aggregated across subjects; *p* < 0.0001). Our network’s classifications were at least as helpful as manual spike-sorting for decoding relative to using threshold crossings (Fig. 6 center; paired, 2-tailed, Wilcoxon signed rank test, Monkey Pe: *p* = 0.75; Monkey Wa: *p* = 0.09). Furthermore, the automated real time operation of our network confers a distinct advantage over manual spike-sorting.

**Figure 6.**
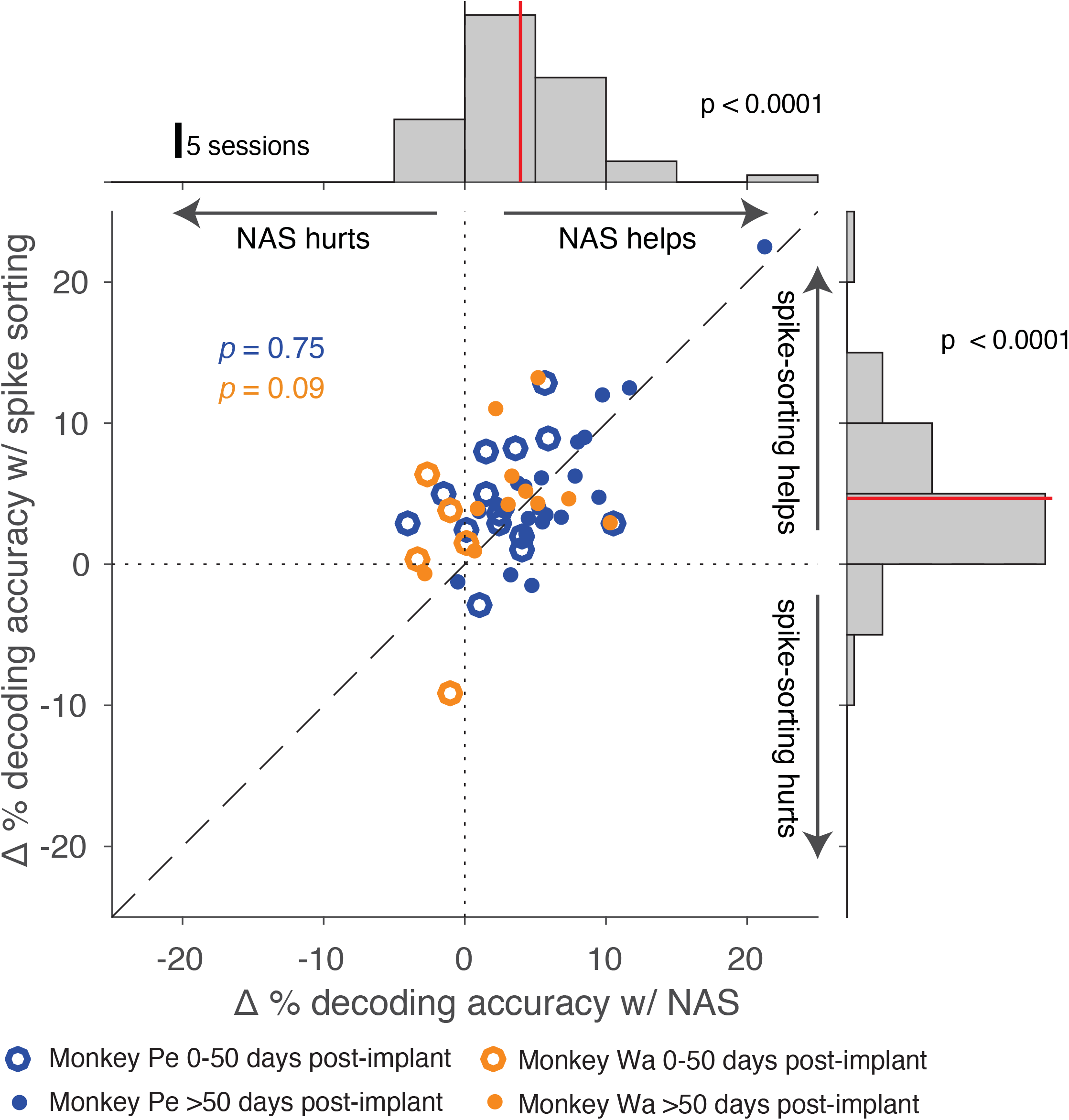
Using NAS classifications was comparable to using spike-sorted data for decoding. Δ % decoding accuracy was calculated for both NAS classifications and spike-sorted waveforms as the change from decoding accuracy with threshold crossings. Top: Distribution of Δ % decoding accuracy using NAS classifications aggregated across subjects (γ threshold selected as in 5d). Using NAS classifications improved decoding accuracy from that with threshold crossings (2-tailed, Wilcoxon signed rank test, p < 0.0001). Right: Distribution of Δ % decoding accuracy using manual spike-sorting aggregated across subjects. Using spike-sorted classifications also improved decoding accuracy from that with threshold crossings (2-tailed, Wilcoxon signed rank test, p < 0.0001). Center: The joint distribution of Δ % decoding accuracy with spike-sorting and NAS classified spikes. Using our network’s classifications was at least as helpful as spike-sorting for decoding (paired, 2-tailed, Wilcoxon signed rank test, Monkey Pe: p = 0.75; Monkey Wa: p = 0.09).

Together, our analyses demonstrate that our network was able to produce classifications of spikes and noise that matched our qualitative expectations and could be tuned quantitatively to adjust its permissiveness. Compared to using threshold crossings, our network’s classifications most often led to an improvement or minimally harmful change in decoding accuracy, indicating that there was usually little downside to the application of our network. These outcomes are valuable particularly in the context of a BCI where real time operation and maximal decoding accuracy are prized.

## Discussion

By leveraging our pool of previously spike-sorted data, we trained a neural network classifier to separate spikes from noise in extracellular electrophysiological recordings in real time. To assess the value of this method, we used an objective criterion – the ability to decode remembered location from prefrontal cortex in a memory-guided saccade task. We found that we could set a fixed γ threshold that resulted in an improvement or at worst a small change in decoding accuracy for the majority of sessions, and this effect persisted even as recording quality decreased over time. Compared to spike-sorting (performed offline with substantial manual labor involved), classifying spikes with our network had similar effects on decoding accuracy relative to threshold crossings and was less tedious.

### Training the network

Developing a neural network trained on spike-sorted data allowed us to capture the essence of the rules by which experienced researchers distinguish spikes from noise without explicitly defining all the variables and exceptions, a standard challenge in spike classification (Wood et al. 2004). Several studies have demonstrated the promise of using a neural network trained on spike-sorted or human-verified data for spike detection and classification (Chandra and Optican 1997; Kim and Kim 2000; Lee et al. 2017; Racz et al. 2019; Saif-Ur-Rehman et al. 2019). Lee *et al*. (2017), Racz et al. (2019), and Saif-Ur-Rehman et al. (2019) used spike-sorted data to train their respective neural networks and developed more complex network architectures than the one-layer network we explored in this work. Each of these studies was validated by the performance of the network in correctly labeling held out spiking waveforms. The intention of our simple architecture was not to compare its performance to existing spike detection and sorting algorithms, but rather to explore how a relatively simple, fast, and trainable noise removal method could improve decoding performance.

We found that a simple neural network was quite good at capturing the nuances of the training set. Experts often disagree on how to classify certain instances of low SNR waveforms. The network was sensitive enough to capture a researcher’s specific spike classification tendencies, such that a network trained on data classified by a particular individual performed better on classifying test data from that individual than from other expert spike classifiers. Given this level of sensitivity, a reasonable concern would be the presence of misclassifications in our training set, as human spike-sorting is both subjective and prone to error (Pedreira et al. 2012; Rossant et al. 2016). However, the network was trained on many waveforms to minimize the impact of any confounders, including waveforms from multiple subjects, brain regions, and human spike-sorters. Additionally, by training the network on labeled data with some variability introduced by human sorters, the network learned to make stronger predictions (higher P(spike) values) for obvious spike waveforms and weaker predictions for more ambiguous waveforms. We then leveraged this range of prediction values to create a tunable parameter for our classifications (i.e. the γ threshold).

An ideal and even more generalizable training set might include waveforms from different research groups, artificially simulated neural waveforms and noise in which ground truth could be established, and/or pre-defined quantities of certain types of spikes (e.g. shifted spikes, slow and fast spike waveforms) and noise (e.g. electrical artifacts) to target what the network learns. However, no matter how much training data is used, an ideal training set could only be defined in the context of objective metrics to assess the trained network’s performance.

### Evaluating the network with decoding

We compared decoding performance with our network classifications to performance with threshold crossings, which are the current standard in the BCI community. We used independent test data to assess the effect of classifications from our trained network on decoding accuracy. Although the expectation might be that removing noise from neural data would improve decoding, historically doing so with spike-sorted classifications has had mixed effects, in some cases hurting performance (Todorova et al. 2014). Nevertheless, many BCIs currently operate with some amount of online noise removal or visual inspection (such as disabling visibly noisy channels) to preprocess neural data before decoding (Chase, Kass, and Schwartz 2012; Hochberg et al. 2006; Homer et al. 2013; Sadtler et al. 2014). The question of whether to spike-sort BCI data prior to decoding or simply use all threshold crossings has been a source of debate with no clear resolution. Our work reframes this question to investigate how an automated noise removal method could be used to aid decoding.

Our findings provide additional support, across multiple sessions over time, that removing certain noise waveforms prior to decoding provides an advantage over using all threshold crossings, consistent with findings from Fraser *et al.* (2009) and Christie *et al.* (2015). Although there were a few occasions when decoding accuracy with threshold crossings was better than decoding using the network, it was never by more than a few percent. Although these few instances might occur merely due to noise in our estimation of decoding accuracy, another possible explanation is that a lack of non-neural noise on the array on certain days would limit the benefits of our network and thereby increase the relative likelihood that waveforms valuable for decoding were removed.

Contrary to the benefits of spike-classification that we observed, a study by Todorova *et al.* (2014) found that using spike-sorted waveforms hurt decoding when noise waveforms were discarded. It is difficult to speculate why removing noise waveforms hurt decoding in their study but helped it in ours given that our methods of noise removal were different. They also found assigning noise waveforms from spike-sorting to a new unit, rather than discarding them, improved decoding relative to threshold crossings. Yet, when we assigned our noise waveforms to separate units, decoding accuracy was worse or unchanged compared to discarding noise waveforms both with our spike-sorted classifications and our network classifications (Supplementary Figure 6, https://doi.org/10.6084/m9.figshare.11808492.v1). Importantly, we defined ‘noise’ operationally as any waveform that did not exceed the γ threshold of our network setting. This definition of noise surely included both electrical artifacts as well as indistinct shapes that were actually neural in origin. Removal of waveforms from the former class could only help decoding, but removal of the latter class might actually harm decoding if the multi-unit activity contained in those waveforms had information about the condition of interest for decoding.

Apparent noise in neuronal activity, in addition to changing the activity of single electrode channels, can also be correlated across channels. In the activity of pairs of single neurons, such fluctuations in trial-to-trial correlated variability are sometimes termed “noise correlation” (also known as spike count correlation, or r_sc_). Noise correlation can have a substantial impact on the ability of a decoder to extract information from neuronal populations (Averbeck, Latham, and Pouget 2006; Kohn et al. 2016), and the activity of multi-unit groups of neurons has a higher noise correlation than that of the constituent pairs of neurons (Cohen and Kohn 2011). Since our network acted on threshold crossings, which typically contain the activity of many individual neurons, it likely also impacted the magnitude of the noise correlation between channels. Our decoder assumed that the noise on each unit was independent, a choice common to BCI, which permitted training the decoder with a smaller quantity of trials than is necessary to learn the covariance structure in the population. The effect of our network on decoding surely depends on not only the structure of the network and the choice of γ threshold, but also the structure of noise present in the population and the sensitivity of the decoder to that noise.

In addition to understanding how the type of noise present in the data impacts decoding, it may also be useful to evaluate the types of neural waveforms that contribute to decoding. Different levels of single and multi-unit activity are captured during a neural recording depending on the threshold set by the researcher (i.e. the minimum voltage level at which the neural activity is marked as a spike). By adjusting this threshold, we change the candidate waveforms available for decoding. Oby *et al.* (2016) found that the optimal threshold depended on the type of information being extracted from neural activity. Another study found that higher thresholds, which captured less multi-unit activity, resulted in worse decoding of direction from M1 (Christie et al. 2015). These studies in the context of our own findings highlight the need to identify specific types of waveforms that contribute to the decodable information in different brain regions and task contexts.

### Other variables that impact decoding accuracy

A key concern for practical implementation of BCIs is decoding stability. Array recordings often get visibly noisier over time, potentially as a result of an immune response to the implant (Ward et al. 2009) or a physical shift in position (Perge et al. 2013). In our study, as the time since array implant increased there were more “low probability” waveforms and fewer “high probability” waveforms, indicating the arrays had fewer well-defined single-unit waveforms. In line with this, several other studies with chronic array implants have found that the number of single-units decreased over time (Dickey et al. 2009; Downey et al. 2018; Fraser and Schwartz 2012; Tolias et al. 2007). Nevertheless, there are mixed findings on the stability of decoding performance in chronic array implants. Some studies have found decoder performance with threshold crossings remained stable over time (Chestek et al. 2011; Flint et al. 2016; Gilja et al. 2012; Nuyujukian et al. 2014); however, other studies, including our own, observed a decrease in decoding accuracy over time (Perge et al. 2014; Wang et al. 2014). Unlike the previous literature investigating the effects of threshold crossings and spike-sorting on decoding performance, we used neural activity from PFC, where decoding accuracy is typically lower (Boulay et al. 2016; Jia et al. 2017; Meyers et al. 2008; Parthasarathy et al. 2017; Rizzuto et al. 2005; Spaak et al. 2017; Tremblay et al. 2015) than in motor cortex (Collinger et al. 2013; Koralek et al. 2012; Masse et al. 2014; Sadtler et al. 2015). Stability of decoding accuracy with array age may also depend on the brain region of the implant, the subject’s task, initial decoding performance, and the choice to train the decoder daily (Chestek et al. 2011; Gilja et al. 2012) or hold it constant (Nuyujukian et al. 2014). The beneficial effects of our network on decoding were particularly salient in the case of older array implants, where decoding accuracy had decreased over time.

### Limitations of NAS

Our network, NAS, is certainly not sufficient to achieve the goal of obtaining well-isolated single-units. NAS evaluates each waveform independently and therefore, cannot distinguish between multiple units recorded on the same channel. A concern for online decoding is that pooling spikes on a given channel could hurt decoding if there are multiple single-units that respond to the task conditions differently. This did not appear to be a major issue in our data because we did not find a substantial difference between decoding performance with pooled spikes from the network classifications and decoding with well-isolated single-units from spike-sorting (Fig. 7).

NAS was trained using previously sorted data available in our laboratory, rather than simulated spike waveforms and noise (Chaure, Rey, and Quian Quiroga 2018) or a ground truth data set (Anastassiou et al. 2015; Neto et al. 2016). This choice had the potential to limit the abilities of NAS to distinguish spikes from noise, as our training data was subject to errors and biases of individual human sorters. However, given the rarity of such ground truth data sets, and the particular features of spikes that are unique to brain areas and the recording hardware in a laboratory, we also view this choice as a strength. Nearly any laboratory performing extracellular electrophysiology would have such training data available, already suitable for their particular brain areas and recording methods. Thus our method is highly customizable for any laboratory, and we view its focus on human-sorted training data as a strength for the particular application to which it was designed.

Additionally, NAS does not take advantage of temporal or spatial information. When manually spike-sorting, sorters often use the inter-spike interval and the stationarity of a candidate unit over the recording to decide whether those waveforms could sensibly belong to a single-unit. For dense arrays and tetrodes, leveraging spatial information is vital for isolating spikes that appear on multiple channels (Chung et al. 2017; Pachitariu et al. 2016). However, such dense recordings are still rare relative to the use of sparse electrodes in electrophysiological experiments. Thus, for traditional offline data processing, NAS could be used as a quick preprocessing step to remove noise and make preliminary spike classifications but must be accompanied by another algorithm or manual spike-sorting to isolate single-units.

### Extensions for NAS

Our neural network-based spike classifier is a promising tool for both offline preprocessing of neural data and improving online decoding performance. However, in designing our network we only scratched the surface of many potential avenues to address these challenges. We found that using networks with different numbers of hidden units and layers did not substantially alter decoding accuracy even though there were some differences in how these different sized networks classified waveforms. Given that our network was relatively simple, it would be possible to implement similar operations with alternative algorithms such as logistic regression. We chose a neural network because it was easily trainable from existing data and there are many ways to modularly build upon its complexity. Although we opted to use 50 hidden units and one hidden layer, it is possible that a more complex network with additional filtering operations and more categories of waveform classification may result in improved decoding performance and could help to create a more robust spike classifier with the ability to distinguish multi-units and single-units for offline analyses. However, our work had the value of exploring how a smaller network trained on spike-sorted data could identify features that were most valuable for decoding.

While training on spike-sorted data from channels with well-isolated single-unit action potentials (SNR>2.5) was an efficient choice because of its easy availability, it may not be the best choice in regimes where canonical spike-shapes do not necessarily carry the most decodable information compared to less well-defined multi-unit activity (Chestek et al. 2011; Stark and Abeles 2007). If the goal is to improve decoding performance, it might be valuable to train a network on waveforms from low SNR channels that contain more multi-unit waveforms or to select waveforms or units that positively contribute to decoding. Ventura and Todorova (2015) developed a method for identifying the information contribution of units for decoding. Such a method could be used to develop a training set of neural waveforms that are assigned a label based on how much they positively or negatively contributed to decoding. Alternatively, if the goal is to discard specific types of artifacts, then it might be ideal to train the network on predominantly low SNR channels that contain those artifacts.

Beyond Bayesian decoding, it would be valuable to assess how NAS classifications impact other neural data analyses. A recent study found that spike-sorting did not provide much benefit over threshold crossings for estimating neural state space trajectories (Trautmann et al. 2019), a powerful demonstration that the principles of neural circuits may be accessible in neuronal population recordings even when single units are not identified. However, this analysis was performed on trial-averaged data. Using NAS classifications might be more helpful in single-trial data where estimating trajectories through neural state space could be negatively influenced by the momentary entry of noise into recordings. Another interesting application would be to assess how NAS classification impacts the measurement of neural noise correlations, which have been shown to be affected by the stringency of spike-sorting (Cohen and Kohn 2011). Using NAS classifications with a low γ threshold could prove to be a useful noise removal tool for applications that are sensitive to classification stringency.

Overall, we developed a new tool for preprocessing BCI data that classified threshold crossings in a tunable manner that was beneficial for decoding. A neural network-based spike classifier has the potential to reduce the need for human intervention in removing noise from neural data. Our tunable classifier is a step toward preprocessing methods that both optimize and stabilize online decoding performance.

## Acknowledgements

We would like to thank Samantha Schmitt for assistance with data collection and Ben Cowley, Byron Yu, and Steve Chase for their valuable insight on this project.

